# SQ3370 Activates Cytotoxic Drug via Click Chemistry at Tumor and Elicits Sustained Responses in Injected & Non-injected Lesions

**DOI:** 10.1101/2020.10.13.337899

**Authors:** S. Srinivasan, N. A. Yee, K. Wu, M. Zakharian, A. Mahmoodi, M. Royzen, J.M. Mejia Oneto

## Abstract

While systemic immuno-oncology therapies have shown remarkable success, only a limited subset of patients benefit from them. Our Click Activated Protodrugs Against Cancer (CAPAC™) Platform is a click chemistry-based approach that activates cancer drugs at a specific tumor with minimal systemic toxicity. CAPAC Platform is agnostic to tumor characteristics that can vary across patients and hence applicable to several types of tumors. We describe the benefits of SQ3370 (lead candidate of CAPAC) to achieve systemic anti-tumor responses in mice bearing two tumors. SQ3370 consists of a biopolymer, injected in a single lesion, followed by systemic doses of an attenuated protodrug of doxorubicin (Dox). SQ3370 was well-tolerated at 5.9-times the maximum dose of conventional Dox, increased survival by 63% and induced a systemic antitumor response against injected and non-injected lesions. The sustained anti-tumor response also correlated with immune activation measured at both lesions. SQ3370 could potentially benefit patients with micro-metastatic lesions.

## Introduction

Recent advances in immunotherapies including checkpoint inhibitors have revolutionized cancer treatments. However, these approaches are limited as they can only treat a subset of patients whose tumors express sufficient levels of necessary biological markers. Additionally, it has been difficult to find biological markers that uniformly predict responses to checkpoint-based therapies. These treatments are often accompanied by several adverse effects including systemic autoimmune reactions that may require hospitalization or long-term treatments. More targeted approaches such as antibody-drug-conjugates (ADCs), nanoparticles, or liposomal formulations have also shown to be efficacious against several cancers. However, these approaches also depend on biological factors, e.g. tumor biomarkers, enzymatic activity, pH, or reactive oxygen levels to distinguish malignant cells from healthy tissues. As these factors are also expressed by healthy cells, albeit at lower levels than tumors, targeted the approaches lack sufficient specificity resulting in off-target toxicities, or are rendered ineffective by resistance mutations thus limiting their clinical potential. The unfortunate outcome is that conventional chemotherapeutic agents continue to be a prevalent component of standard treatment for a wide variety of tumors despite their narrow therapeutic window and debilitating systemic toxicities.

We now present Click Activated Protodrugs Against Cancer (CAPAC), a modular approach that enables the activation of cancer drugs at a specific site in the body with minimal systemic toxicity. Click chemistry is a class of chemical reactions between mutually reactive agents that occur with high efficiency and specificity, without the need for special solvents or catalysts^1^. The reaction between tetrazines and *trans-cyclooctene* (TCO) is one of the most efficient click reactions reported to date, with demonstrated utility in animal models systems.^2–9^ Tetrazine and TCO react irreversibly and specifically *in vivo* with a high thermodynamic driving force, with minimal interaction with the surrounding biological mileu.^10,11^ The CAPAC Platform uses a tetrazine-modified biopolymer injected at the tumor that can capture a systemically circulating TCO-modified protodrug™, to precisely release the active drug at the tumor. (**Figure 1a**). We define a protodrug as a chemically modified form of the active or parent drug that has attenuated toxicity and activity compared to the active drug. The protodrug can be converted into the active drug only upon interaction with the biopolymer in the CAPAC platform. Thus, the platform does not rely on innate differences in biochemistry or tumor biomarkers to achieve tumor specificity. Rather, it depends on the chemical properties of the tetrazine/TCO reaction. This increased selectivity results in a wider therapeutic window compared to other targeted approaches. Additionally, because the 2 components are administered separately, the CAPAC Platform allows for repeated dosing of the intravenous protodrug with a single local injection of the biopolymer, thereby offering temporal control.

**Figure 1:**
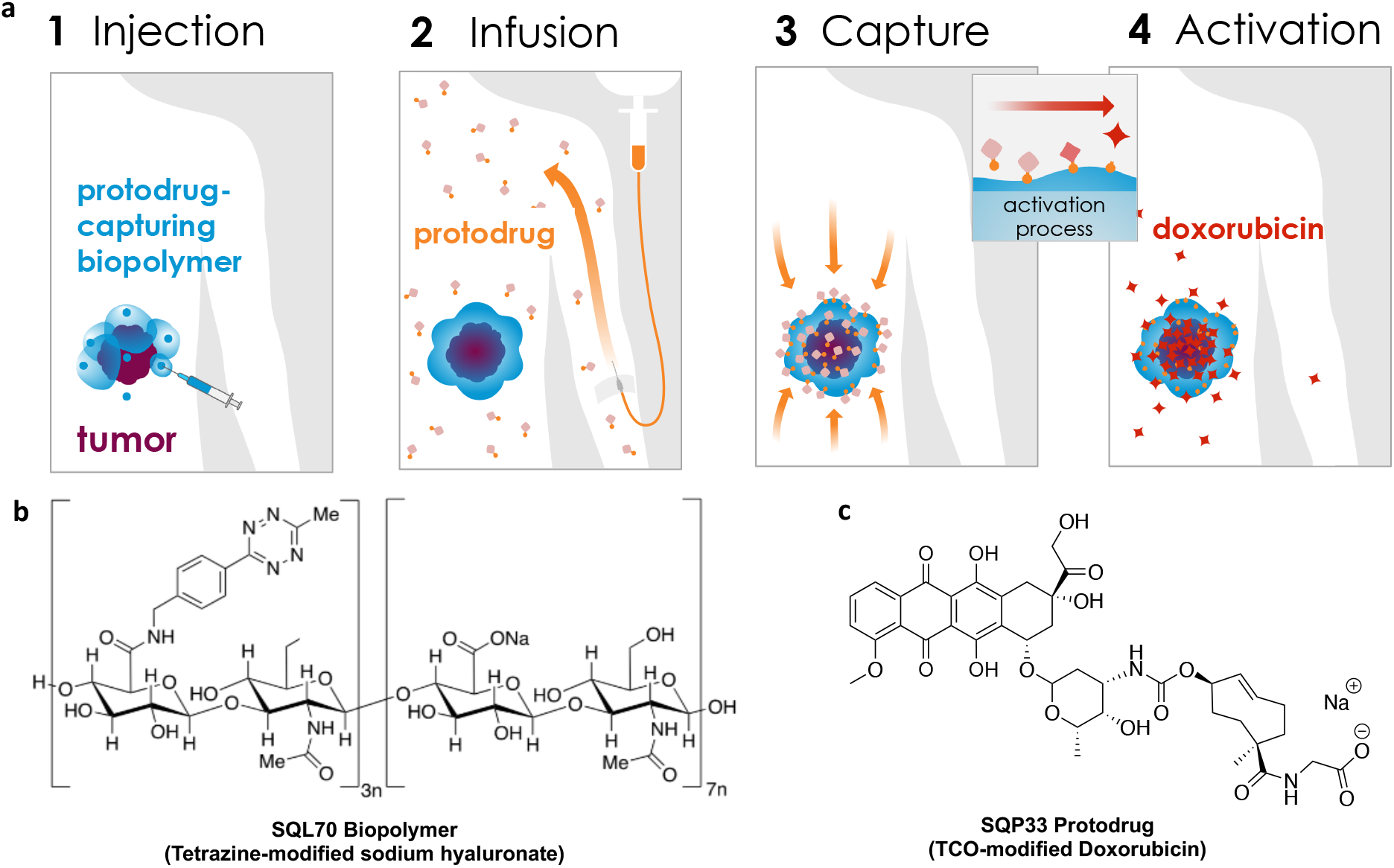
Platform Mechanism and SQ3370 Components. a) SQL70 biopolymer is locally injected at the tumor site, and SQP33 protodrug is infused systemically. SQP33 protodrug is captured by SQL70 biopolymer at the tumor site through a rapid covalent reaction between tetrazine (blue) and *trans-cyclooctene* (TCO; orange) moieties, followed by chemical rearrangement to release active Dox. b) SQL70 biopolymer is a tetrazine-modified form of sodium hyaluronate. c) SQP33 protodrug consists of doxorubicin conjugated to a reactive TCO group.

In 2014, we demonstrated the proof-of-concept and feasibility of the technology in an animal model.^2^ In 2016, we showed the enhanced safety and sustained anti-tumor responses using this technology in a mouse fibrosarcoma model with a single tumor^3^. Here, we discuss the effects of SQ3370, the lead asset of the CAPAC Platform, in a dual-tumor mouse model. SQ**3370**, consists of SQL**70**, a hyaluronic acid-based biopolymer that is modified with tetrazine (**Figure 1b**), and SQP**33**, a protodrug of doxorubicin (Dox) that is modified with TCO (**Figure 1c**). An Investigational New Drug (IND) application filed for SQ3370, received regulatory clearance for Phase I clinical testing in patients. Hyaluronic acid (HA) was chosen as the backbone of SQL70 biopolymer over alginate^3^ due to its excellent tolerability and its function as a regulator of inflammation.^12^ Hyaluronic acid is a ubiquitous, non-sulfated glycosaminoglycan that is well tolerated in the body, and has been used clinically in unmodified, crosslinked, and derivatized form for over five decades.^13,14^ Extensive process development including manufacturing scale-up was done to optimize tetrazine conjugation, pH, purity, viscosity and injectability of SQL70 biopolymer. Similarly, the SQP33 protodrug was optimized for improved aqueous solubility, stability, manufacturability, purity and increased yield. Details on the process optimization of both components will be published separately. SQ3370 is being tested in patients with advanced solid tumors for whom first-line treatment has failed. SQ3370-001, the single-arm, open-label first-inhuman Phase I clinical trial is open and enrolling in the United States and Australia (clinicaltrials.gov/ct2/show/NCT04106492).

Sustained tumor responses are often sought after in cancer treatments but are hard to achieve in the clinic.^15–17^ Long-term remission usually depends on underlying mechanisms that activate the patient’s own immune system to combat tumors.^17–19^ One such mechanism to trigger immune activation is through the recognition of damage-associated molecular patterns by antigen-presenting cells that later prime T cells against tumor-associated antigens^20^. Certain chemotherapies, including Dox, are known to induce immune activation that triggers an immune response against tumors^21^. However, prominent systemic side effects of Dox such as severe lymphopenia and cardiotoxicity limit the amount that can be given and its potential benefits. We hypothesized that a Dox-based treatment combining a high local therapeutic index with low systemic toxicities may enhance the intrinsic effects of Dox, and in turn produce an anti-tumor effect that could treat both local and metastatic lesions. Accordingly, using an immune adjuvant, such as a toll-like receptor (TLR) agonist, may further enhance the immune effect of Dox. TLR agonists have shown strong anti-tumor effects, either as monotherapy or in combinations.^22^ While no systemic TLR agonists are clinically approved to date, three have been approved for local injection in cancer treatment and many others are being evaluated in pre-clinical and clinical studies.^23^ When injected into the primary tumor, TLR agonists trigger an immune response that causes tumor regression at distant sites in mouse models of breast carcinoma, colon cancer, melanoma, and lymphoma, as well as spontaneous tumors.^24,25^ In advanced colorectal cancer, for example, patient survival can be prolonged if metastatic lesions can be successfully treated, usually by surgical resection of all gross disease.^26,27^ While non-invasive treatment options are available for patients with oligometastatic disease who are ineligible for surgery, they are usually less effective.^28^ For these patients, combining TLR agonists with chemotherapy could improve the latter’s efficacy against metastatic tumors.^29,30^

Here, in a colorectal carcinoma MC38 murine model with two subcutaneous tumors, we explore the ability of local activation of Dox using SQ3370 with or without a TLR agonist to induce systemic anti-tumor responses.^31,32^

## Results

### Biopolymer Selection and Improving Protodrug Physical Properties

The choice of HA as the biopolymer backbone offers a highly biocompatible foundation^12^ with tunable material properties to create a product suitable for administration in patients. Hence, tetrazine-modified HA was selected for further optimization, replacing the previous alginate-based prototype.^3^ After evaluating several HA molecular weights, molar ratios of reagents, and degrees of substitution, we ascertained a tetrazine-modified HA formulation that was straightforward to manufacture (unpublished raw data). This material also offered sufficient fluidity to enable peritumoral injection. We named this material, SQL70 biopolymer(**Figure 1b**).

Our initial studies^3^ using the first-generation protodrug, TCO-Dox, showed that although supratherapeutic doses of Dox could be activated locally with the technology platform, the first generation protodrug dose was still limited by its low aqueous solubility. To improve the solubility of the protodrug, we introduced a hydrophilic group to the TCO moiety. The second generation of this protodrug, SQP33, contains a glycine residue (**Figure 1c**), and was soluble in pH 7 phosphate-buffered saline (PBS) at 72 mg/mL, an increase of 4 orders of magnitude over the first-generation TCO-Dox protodrug. Moreover, studies that measured the extent of attenuation of SQP33’s cytotoxicity revealed that SQP33 was 82-fold less cytotoxic towards MC38 cancer cells than Dox (**Figure S1**). This translated to higher tolerability *in vivo* as well, where mice received a cumulative dose of SQP33 that was >5.9-fold higher than the maximum dose of conventional Dox, with no significant observable side effects including drop in body weights (**Figure 2, S2**).

**Figure 2:**
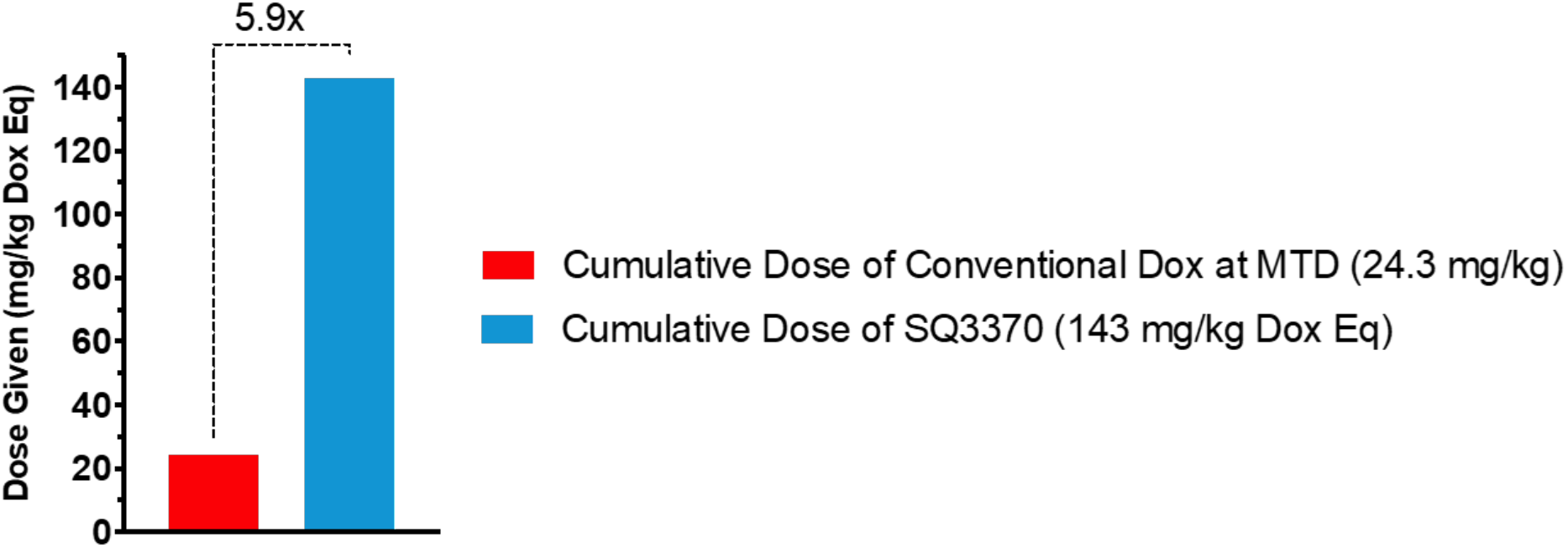
Dose Comparison Between Multiple-Dose SQ3370 and Doxorubicin Hydrochloride in Mice. C57BL/6 were treated intravenously with SQ3370 (5 doses of SQP33 protodrug following a single subcutaneous injection of SQL70 biopolymer). Animals given only Dox HCl at the 8.1 mg/kg Q4D x 3 days were used as a control. When given over 5 cycles (28.6 mg/kg QD Dox eq x 5 days), SQP33 was well tolerated at 5.9 times the dose of conventional Dox.

#### Anti-tumor effects of SQ3370 in immune-competent mice with dual MC38 tumors

Meeting our objectives for obtaining suitable physical properties of SQL70 biopolymer and SQP33 protodrug, and increased tolerability in mice, led us to evaluate the anti-tumor effects of the SQ3370 in a syngeneic model in immunocompetent mice with MC38 tumors. Mice were subcutaneously inoculated with two tumors — a large primary tumor, also called the “injected tumor” as it was later injected with SQL70 biopolymer at the start of treatment, and a small distal tumor, called the “non-injected tumor,” as it was not injected with SQL70 biopolymer (**Figure 3a**). While this model is not representative of true metastasis, it was used as a surrogate to assess the effect of SQ3370 on a distal tumor. We also assessed the effects of the treatment on the overall survival, local and systemic anti-tumor responses.

**Figure 3:**
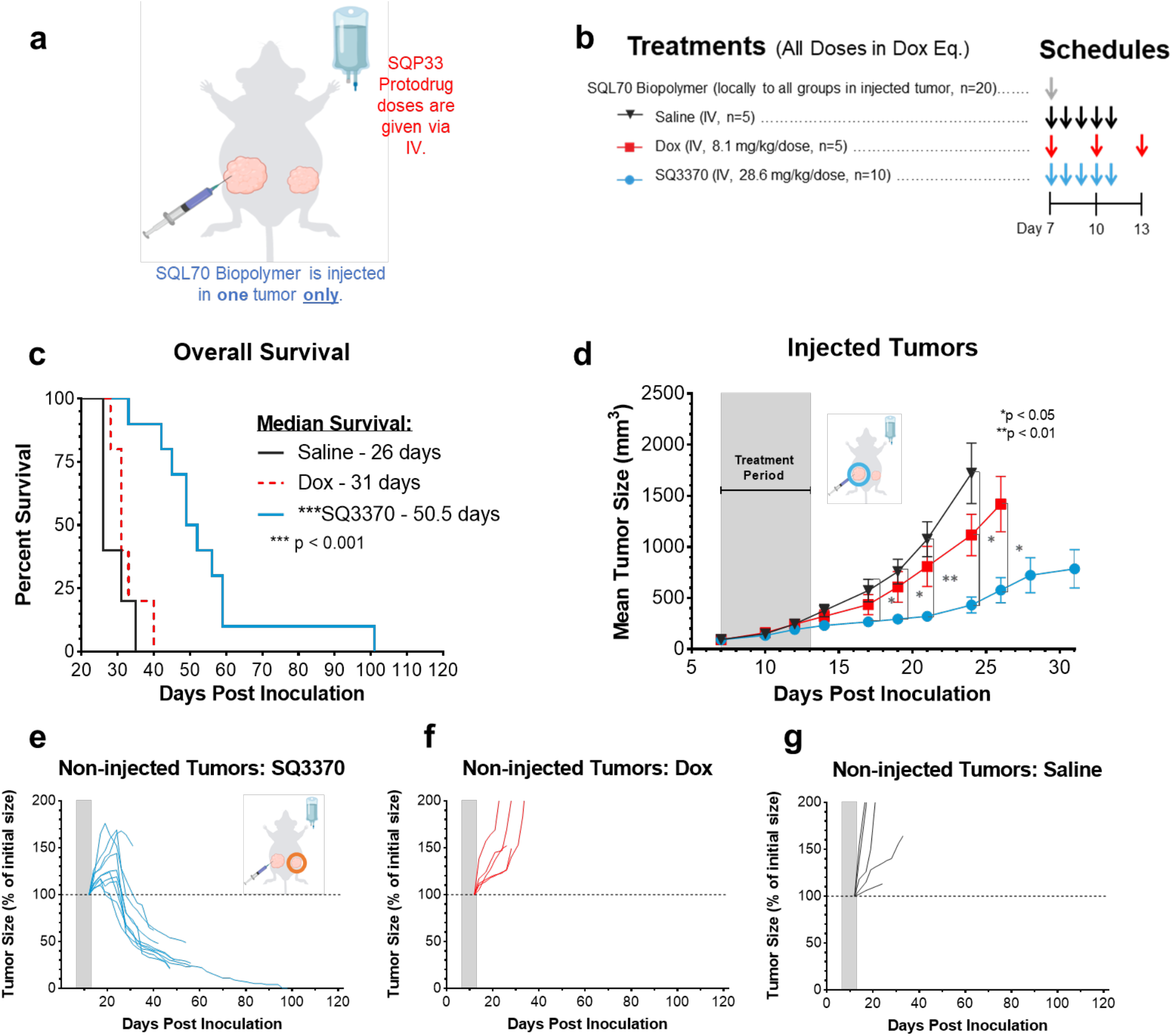
SQ3370 treatment — Tumor Response and Overall Survival in Dual-Tumor Mice. a) Treatment schematic for a syngeneic MC38 dual tumor study in immunocompetent C56BL/6 mice. All tumor cells were implanted at Day 0. b) Treatments started at Day 7 with local injection of biopolymer at the primary “injected” tumor, followed by systemic therapies. c) Kaplan-Meier survival curves for treatment groups. d) Tumor growth curves show mean ± SEM of large primary tumors injected with SQL70 biopolymer (injected tumors). e to g) Spider plots showing the growth of individual distal non-injected tumors in SQ3370, Dox, and saline treated groups, respectively. Tumor growth curves of individual non-injected tumors are displayed as a percentage of the initial volume of each tumor (measurement from day 12 post-inoculation) for each treatment group. Data points without errors bars occurred when the standard error was smaller than the symbol used to represent the treatment condition. Curves stopped after 1 or more mice in that group died or were sacrificed when tumor volume reached 2000 mm3. Gray bars represent treatment duration. Statistical significance in tumor growth curves was determined by unpaired t-test with Welch’s correction for each day. Statistical significance in survival was determined by log-rank (Mantel-Cox) test. *P<0.05; **P<0.01;***P<0.001.

SQ3370 (**Figure 3b**) resulted in an improved median survival of 50.5 days which was 63% better than control Dox treatment (31 days) or 94% better than saline (26 days) (**Figure 3c**, p < 0.001). SQ3370 also produced a sustained anti-tumor response of the injected tumors (**Figure 3d, p<0.05**) compared to control Dox or saline treatment. Interestingly, SQ3370 treatment in mice resulted in tumor regression of the non-injected tumors (**Figure 3e**), while Dox or saline did not (**Figure 3f,g**). Per our hypothesis, we observed that activating Dox locally with SQ3370 in contrast to systemic delivery, produced a sustained anti-tumor effect against injected and non-injected lesions.

To determine if the sustained anti-tumor response correlated with immune activation, we performed multi-color flow cytometric analysis on tumors 2 weeks after SQ3370 treatment (**Figure 4a**). We assessed the biomarkers of tumor-infiltrating lymphocytes (TILs); the corresponding cell populations of interest are provided in **Table 1**. SQ3370 increased the total percentage of TILs or infiltrating T cells (CD45+ CD3+) in both injected (**Figure 4b**) and non-injected tumors (**Figure 4c**) compared to saline. Of these TILs, we observed a higher percentage of CD4+ T cells (helper T cells) in both lesions (p<0.05). This was accompanied by increased percentage of CD8+ T cells (cytotoxic T cells) in injected tumors (p<0.05) and decreased CD4+ CD25+ FoxP3+ T cells (regulatory T cells or T_reg_s) in non-injected tumors with SQ3370 treatment (p <0.01). SQ3370 also increased PD-1 expression among CD4+ T cells in the injected tumors alone. Collectively, the biomarker analysis suggested an increase in the immune activation against the tumor in both injected and non-injected tumors after 2 weeks when treated with SQ3370. This result correlates with sustained anti-tumor responses seen in **Figure 3**, further supporting our hypothesis that local Dox activation at the injected tumor site induces immune activation.

**Figure 4:**
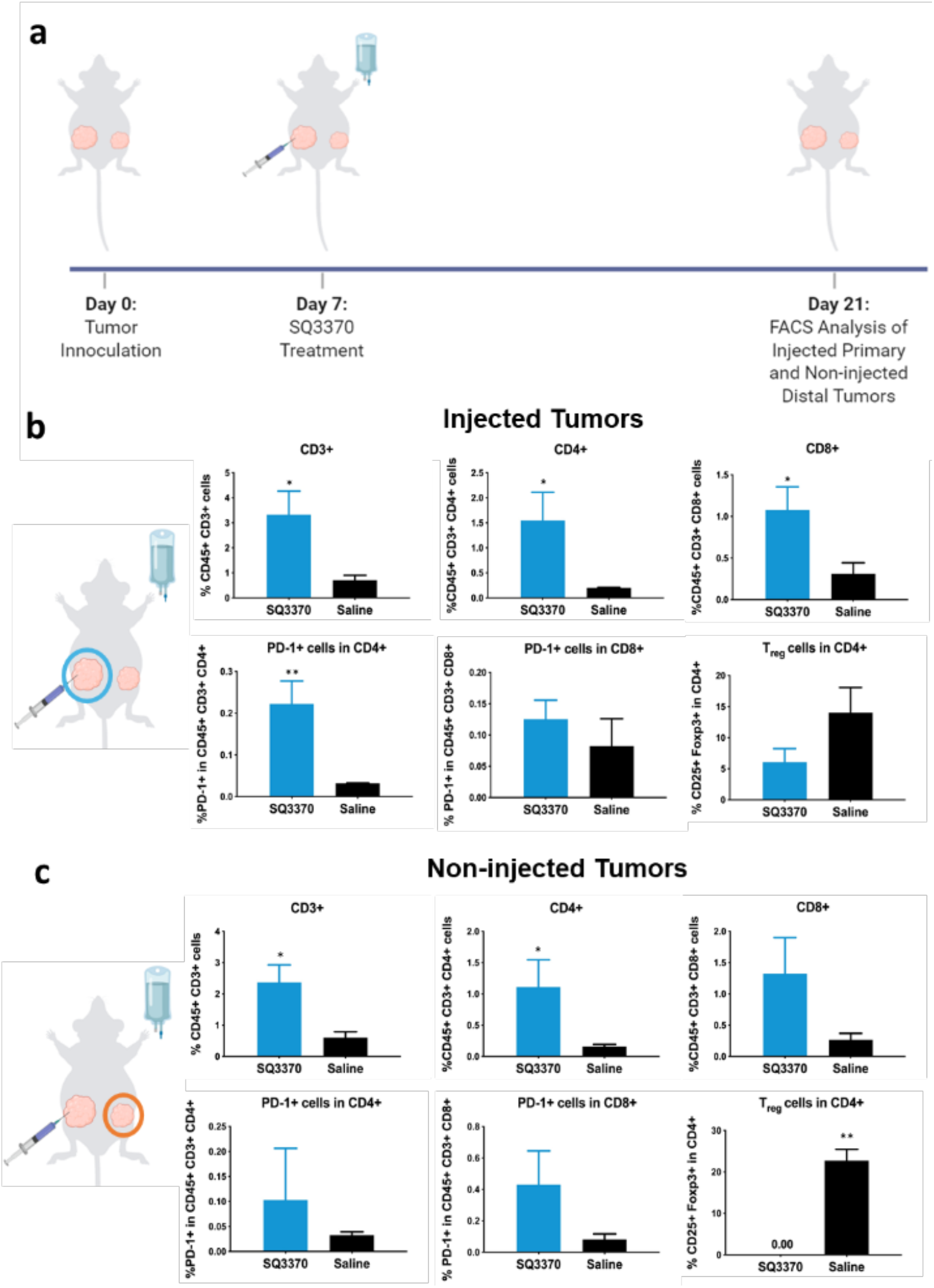
Tumor-infiltrating immune cell profile at 2 weeks post SQ3370 treatment. a) Primary injected and distal non-injected MC38 tumors were harvested from SQ3370 and saline treated mice and analyzed by flow cytometry 2 weeks after treatment. The b) primary injected tumor and c) distal non-injected tumor samples were stained with a cocktail of antibodies against several markers including CD45, CD3, CD4, CD8, PD-1 and FoxP3. The samples were analyzed by multicolor flow cytometry. Dead cells were excluded from analysis. Figure shows the Mean ± SEM of n=5 as per group of multi-stained cells as a percentage of total cells obtained from the tumor sample. Statistical analysis was done by comparing the means of the SQ3370 group with the saline group using a corrected Student’s t-test. \**P*<0.05; \*\**P*<0.01

**Table 1:** FACS Markers and Target Cell Populations

| Biomarkers | Cell populations |
| --- | --- |
| CD45+ | Infiltrating immune cells in the tumor |
| CD45+ CD3+ | T cells in infiltrating immune cells |
| CD45+ CD3+ CD4+ | CD4+ T cells in total T cells |
| CD45+ CD3+ CD8+ | CD8+ T cells in total T cells |
| PD-1+ cells in CD45+ CD3+ CD4+ | Exhausted cells in total CD4+ T cells |
| PD-1+ cells in CD45+ CD3+ CD8+ | Exhausted cells in total CD8+ T cells |
| CD25+ Foxp3+ in CD45+ CD3+ CD4+ | T<sub>reg</sub>s in total CD4+ T cells |

#### Effects of Combining SQ3370 with an Immune Adjuvant

We also evaluated if the anti-tumor effects of SQ3370 could be further enhanced by adding an immune adjuvant, TLR9a. As before, mice with two MC38 tumors each were treated with SQ3370 + TLR9a and compared to SQ3370 alone or Dox + TLR9a control (**Figure 5a,b**). Only the injected tumors received the TLR9a intratumorally. TLR9a addition to SQ3370 prolonged the median survival (101 days) compared to SQ3370 alone (50.5 days) or Dox + TLR9a control (42 days) (p<0.01, **Figure 5c**). TLR9a with SQ3370 further improved the sustained anti-tumor response in injected tumors compared to SQ3370 treatment alone (p<0.05 **Figure 5d**). SQ3370 + TLR9a or SQ3370 alone resulted in robust and sustained tumor regression of non-injected tumors compared to Dox+TLR9 control-treated mice (**Figure 5e-g**). Further, SQ3370+TLR9 induced complete remission of the non-injected tumors (8 out of 10 mice) compared to the SQ3370-treated group (1 out of 10 mice). Therefore, combining TLR9a treatment with SQ3370 improved overall disease outcomes of anti-tumor efficacy, and survival. On the other hand, combining TLR9a with conventional Dox control did not elicit anti-tumor effects on the injected and noninjected tumors, suggesting that there likely is a benefit due to activating Dox locally with SQ3370 in combination with the intratumoral TLR9a. This result, at least in part, may be dependent on TLR9a’s ability to heighten the immune activation induced by SQ3370.

To further assess the presence of a systemic anti-tumor response, complete responders from the SQ3370 alone and SQ3370 + TLR9a treatment groups were re-inoculated with MC38 cells (same number of cells as the primary tumor inoculation, 5 x 10^5^ cells). In these animals, the growth of MC38 tumors upon re-challenge was completely suppressed without the need for additional treatments (**Figure S3**).

**Figure 5.**
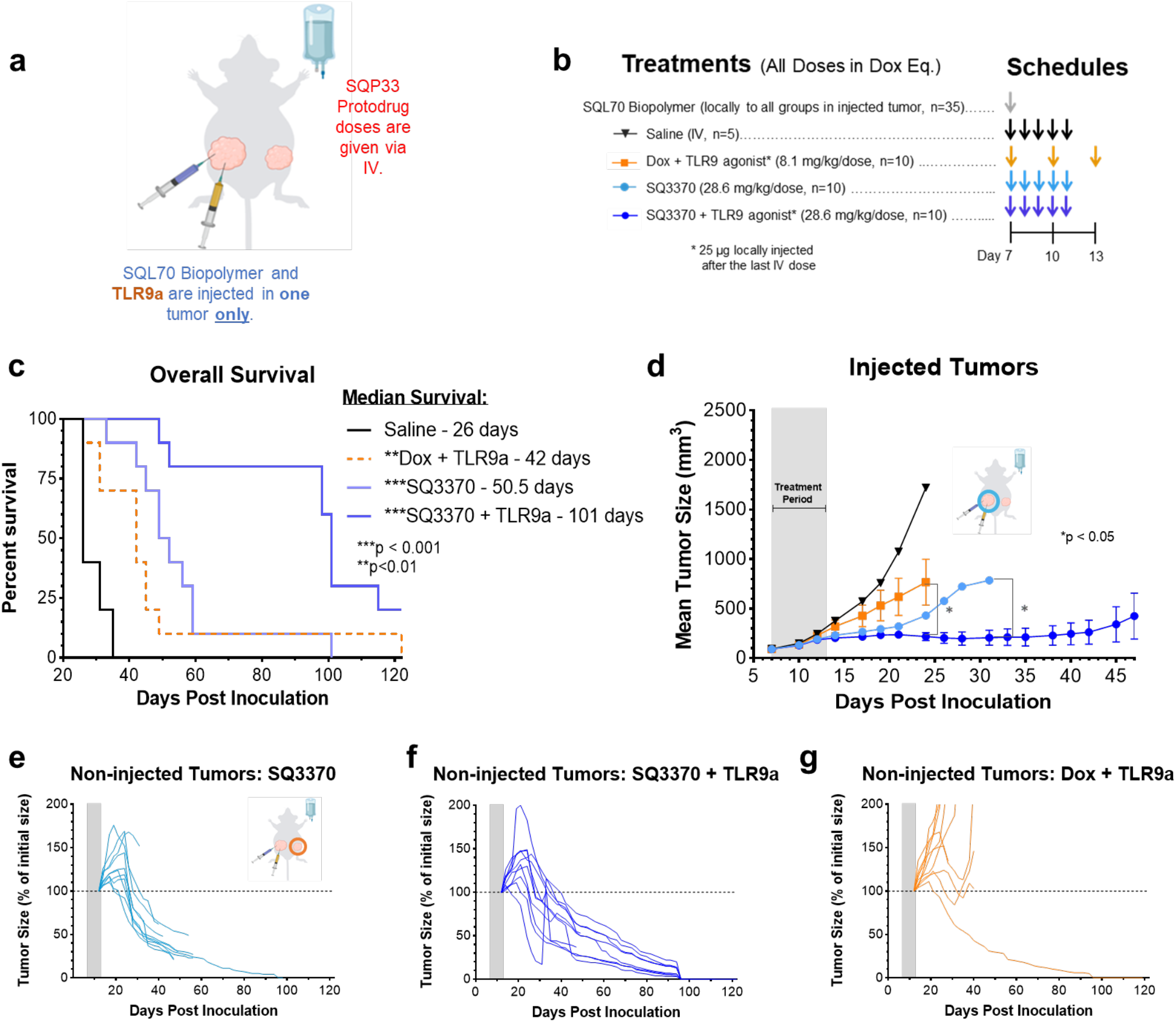
SQ3370 treatment with TLR9a— Tumor Growth and Overall Survival in Dual-Tumor Mice. a) Treatment schematic for a syngeneic MC38 dual tumor study in immunocompetent C56BL/6 mice. All tumor cells were implanted at Day 0. b) Treatments started at Day 7 with local injection of biopolymer at the primary “injected” tumor, followed by systemic therapies and a local injection of 25 ug TLR9a at the “injected” tumor after the last IV dose. c) Kaplan-Meier survival curves for animals treated with Saline, Dox with TLR9a, SQ3370, or SQ3370 with TLR9a. d) Tumor growth curves show mean ± SEM of large primary tumors injected with SQL70 biopolymer (injected tumors). e to g) Spider plots showing the growth of individual distal non-injected tumors in SQ3370, SQ3370 with TLR9a, and Dox with TLR9a treated groups, respectively. Tumor growth curves of individual non-injected tumors are displayed as a percentage of the initial volume of each tumor (measurement from day 12 post-inoculation) for each treatment group. Data points without errors bars occurred when the standard error was smaller than the symbol used to represent the treatment condition. Curves stopped after 1 or more mice in that group died or were sacrificed when tumor volume reached 2000 mm3. Gray bars represent treatment duration. Statistical significance in tumor growth curves was determined by unpaired t-test with Welch’s correction for each day. Statistical significance in survival was determined by log-rank (Mantel-Cox) test. *P<0.05; **P<0.01; ***P<0.001.

## Discussion

In 2016, we demonstrated the safety and efficacy of the CAPAC Platform in a single-tumor model of mouse fibrosarcoma.^3^ We then developed SQ3370, the lead candidate of the platform, in which several modifications were made to the physical properties of its components SQL70 biopolymer and SQP33 protodrug, to optimize them for human clinical use.

Here, we have investigated the effects of SQ3370 in a dual tumor model of colorectal cancer (MC38) in immunocompetent animals in which only the larger tumor was injected with the biopolymer (injected tumor), representing a target lesion receiving the full benefit of local cytotoxic activation. The smaller distal tumor, representative of a metastatic lesion, did not receive any biopolymer injection and hence, experienced no local cytotoxic activation. Eradication of the smaller tumor was only reliant on a systemic anti-tumor effect. In this model, SQ3370 effectively suppressed both the injected and non-injected tumors, and increased median survival over Dox treatment (**Figure 3**). SQ3370 treatment increased TIL biomarkers in injected and non-injected tumors compared to saline treatment. Key differences in TIL subsets were noted. Namely, a significant decrease in FoxP3+ T_reg_s was only seen in the non-injected distal tumor (**Figure 4c**), while a significant increase in CD8+ T-cells was only seen in the injected primary tumor (**Figure 4b**). Both the increase in CD8+ T cells and decrease in T_reg_s are characteristic of an immune activation response albeit by different mechanisms.^33^ The inherent differences in the tumor sizes or local activation of therapy may have caused variations in the infiltrating T-cell subsets. Elucidation of the mechanism behind these differences is the target of ongoing investigations.

Activation of the immune system against tumors appears crucial for producing a long-term sustained anti-tumor response.^34^ Although most chemotherapies cause immunosuppression, certain agents, including Dox, can trigger immune activation against the tumor.^21^ Systemic treatment with conventional Dox has a narrow therapeutic window for triggering this phenomenon. In contrast, we hypothesized that local activation of Dox with SQ3370 directly at the tumor would broaden its therapeutic window by significantly reducing systemic toxicity and potentially trigger the immune activation by inducing immunogenicity of dying tumor cells. Anti-tumor effects combined with a heightened immune response seen in the dual-tumor model after SQ3370 treatment corroborate this hypothesis.

In addition to causing immune activation, SQ3370 treatment resulted in an increase in PD-1 expression in TIL subsets (**Figure 4**). PD-1 upregulation indicates T-cell exhaustion and a potential loss of T-cell effector functions.^35,36^ PD-1 can be induced by the tumor in an attempt to overcome the immune system’s attack.^37^ In our studies, PD-1 expression was observed in the larger, injected tumor but not in the smaller non-injected tumor likely due to the inherent size differences between the two. It may have also been due to the differences in Dox activation the two different locations. Nevertheless, PD-1 expression in TILs upon SQ3370 treatment makes the latter a promising candidate for combination therapies with anti-PD-1 checkpoint blockers in future studies. Chemotherapy-induced immune activation has been shown to sensitize tumors to the cytotoxic function of CTLs^38^ and immune checkpoint blockade therapies.^39^ Addition of Dox-based therapies to anti-PD-1 and anti-CTLA-4 monoclonal antibodies (mAbs) has resulted in significantly improved anti-tumor response compared with the antibodies alone^40^. Overall, combinatorial strategies with locally activated Dox may be beneficial in treating “cold” tumors by making them immunogenic and susceptible to checkpoint inhibition-based immunotherapies.

TLR agonists are known immune adjuvants that enhance the effects of immune activation via antigen presentation^41^ making them another option to explore for combination treatment with SQ3370. Here, a commercially available TLR9a was chosen based on recent studies that demonstrated that intratumoral delivery of the compound triggered a T-cell-mediated anti-tumor response that suppressed local and distal tumors.^24,25^ Combination of SQ3370 with TLR9a doubled overall survival, improved the anti-tumor response and increased the number of complete responders compared to SQ3370 alone. TLR9a was able to heighten the anti-tumor efficacy of SQ3370 likely by enhancing the latter’s immune activation effects, thus, providing further support to our local Dox-induced immune activation hypothesis.

The differences between the SQ3370 and conventional Dox treatment groups highlight that local Dox activation may significantly enhance the systemic anti-tumor response. Compared to conventional Dox, SQ3370 treatment resulted in an improved anti-tumor response. The same trend was observed when each agent was given in combination with TLR9a. Notably, SQ3370 treatment without an immune adjuvant achieved greater anti-tumor efficacy than the combination of conventional Dox and TLR9a – in particular, resulting in a greater number of complete responses in the non-injected tumors. Thus, SQ3370 treatment alone was sufficient to induce a systemic anti-tumor response, even in the absence of an additional immune adjuvant. In combination with a TLR9a, the effects of SQ3370 were even further pronounced. We expect that an immune-mediated anti-tumor effect may be enhanced when the locally activated therapy precludes systemic immunosuppression, thus enabling the CAPAC Platform to achieve systemic responses which were not possible with conventional therapy. We propose a potential mechanism of action for the local and systemic anti-tumor effects of SQ3370 in **Figure 6**.

**Figure 4:**
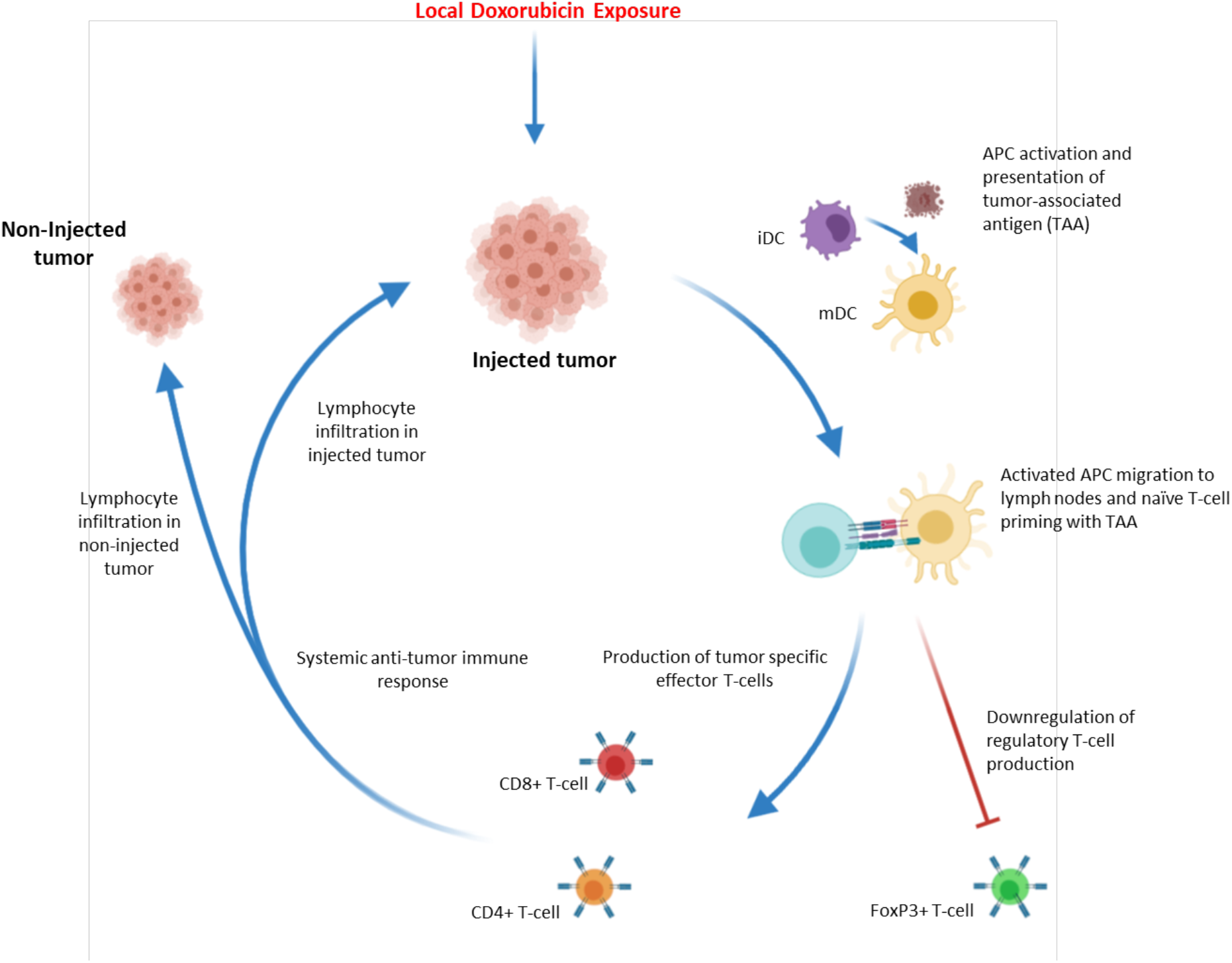
SQ3370 Local Drug Activation Technology triggers systemic anti-tumor immune response. Local Dox activation and exposure in injected tumor site activates APC cells such as iDCs (immature dendritic cells). Activated cells, now mDCs (mature dendritic cells), migrate to lymph nodes and present tumor associated antigen (TAA) to naïve t-cells. The interaction leads to expansion of tumor specific effector T-cells and down regulation of tumor specific regulatory T-cells. Effector T-cells then move to the periphery and generate a systemic anti-tumor immune response, leading to lymphocyte infiltration into both injected and non-injected tumors.

The CAPAC Platform addresses the shortcomings of current immunotherapy and targeted treatments that rely on the inherent differences in biochemistry between healthy and malignant cells. First, the platform is agnostic to inherent differences in the tumor biochemistry that can vary from patient to patient. Instead, its specificity is dependent on the properties of tetrazine/TCO reaction. With SQ3370, a single injection of SQL70 biopolymer at the tumor can activate multiple intravenous doses of SQP33 protodrug with the expectation of reduced acute and cumulative toxicity. Second, with this platform, a higher therapeutic intratumoral concentration may be achieved than would have been possible with conventional chemotherapy, resulting in more tumor destruction.

The ability to reach higher concentrations of cytotoxic therapy in tumors without observable systemic side effects may allow the CAPAC Platform to be used in many potential scenarios. For example, it could be used in cancer treatments that rely on systemic cytotoxic agents as first-line treatment and may also treat cancers that do not respond to immunotherapy or other targeted treatments. More specifically, it could be used in patients with localized tumors that are accessible by local injections but are unresectable due to the size or location of the tumors, such as in head and neck cancers. The platform could also be used in metastatic or disseminated disease in which at least one lesion is accessible by local injection such as in metastatic melanoma, colorectal or breast cancers. It would also be an ideal therapeutic candidate where conventional chemotherapy carries too high of a risk for practical use such as in neoadjuvant therapy, symptomatic tumors ineligible for other treatment modalities (such as previously irradiated disease), or when a patient is too weak for conventional chemotherapy. The platform could even be applied in post-resection adjuvant therapy, in which the biopolymer could be injected directly into the resection cavity in the absence of the tumor. From our preclinical studies, we’ve shown that the biopolymer can be effective at activating protodrug when it is injected in a variety of locations without the presence of a tumor, such subcutaneous,^4^ intramuscular, intra-articular or intraperitoneal regions (unpublished raw data). Further, SQ3370 could be used to convert “cold tumors” to “hot tumors” by making them more immunogenic and responsive to checkpoint inhibitor-based immunotherapies.

## Conclusion

In summary, the CAPAC platform is capable of locally activating potent therapy at the tumor site. SQ3370, the lead candidate of the CAPAC Platform, elicited sustained local and systemic anti-tumor responses that were accompanied by systemic immune activation in a dual tumor model in immunocompetent mice. By locally activating Dox with SQ3370, we likely expanded its pharmacological capabilities while minimizing its systemic toxicity. Moreover, combining SQ3370 with an immune adjuvant, TLR9a, boosted the overall anti-tumor efficacy, by potentially enhancing SQ3370’s immune activation effects. SQ3370 could be used as a treatment option for a wide variety of local and disseminated tumors in the clinic. In the near future, we look forward to reporting results from the Phase I clinical trial in which SQ3370 is being tested in patients with advanced solid tumors.

## Methods

### MC38 Tumor model and flow cytometry

Immunocompetent male C57BL/6 mice were inoculated SC in the right flank with 5 x 10^5^ MC38 tumor cells (large injected tumor) and in the left flank with 1 x 10^5^ MC38 tumor cells (small noninjected tumor). Prior to injection, the MC38 cells were suspended in 0.1 mL DMEM media mixed with 50% Matrigel for tumor development. When the average tumor volume of the large tumor reached approximately 100 mm^3^, animals were randomly grouped according to body weight and tumor volume into 5 treatment groups with 5-10 mice per group. All 5 groups received peritumoral biopolymer injections (100 μL/mouse) near the large tumor (hereafter, the injected tumor). One hour later, groups 1 and were IV administered saline control (QD x 5 days); groups 3 and 4 were IV administered SQP33 protodrug (28.6 mg/kg Dox Eq QD x 5 days; cumulative dose of 143 mg/kg Dox Eq) and group 5 was IV administered Dox HCl control (8.1 mg/kg Q4D x 3 doses; cumulative dose of 24.3 mg/kg). Tumor volumes for both tumors were measured three times weekly in two dimensions using a caliper, and the volume expressed in mm^3^. Three groups, 1, 3 and 5 (saline, n=5; Dox HCl, n=5, SQP33, n=10) were used to assess the tumor growth inhibition, while groups 2 and 4 (saline, SQP33 n=10/group) were used to determine the immune cell infiltration using flow cytometry. Additional 2 groups (n=10) were used to test SQ3370 + TLR9a (SL-01) and Dox + TLR9a. The TLR9a was given as an intratumoral injection in the primary tumors alone after the last SQP33 or Dox dose at 25 μg per mouse. Flow cytometric analysis was carried out with the animal groups 2 and 4 (saline, SQP33 n=10/group) to determine the immune cell infiltration. Tumor collection was performed on the second and fourth subgroups at 1 and 2 weeks, respectively. Followed by RBC lysis and Fc-blocking, the tumor-derived cells were analyzed for cell-surface (CD45, CD3, CD4, CD8, CD25, PD-1) or intracellular (FoxP3) markers using flow cytometry. Cells were also marked with a live/dead stain (Fixable Viability Stain) to discriminate non-viable from viable cells. CD45 is a cell-surface (transmembrane) molecule available on most cells of hematopoietic origin (i.e. from blood). It is used to distinguish infiltrated cells from native cells of the MC38 tumor, which is a colon carcinoma line and lacks CD45. CD3 is a pan T-cell marker, while CD4 and CD8 are present on Helper T cell (T_H_ cells) and cytotoxic T lymphocyte (CTL) subsets, respectively. FoxP3 is an intracellular marker found on CD4+ CD25+ cells and are typically identified as the most common type of T_reg_s. PD-1 is an immune-checkpoint protein and a programmed cell-death receptor found on cells (commonly T-cells). When bound by its cognate ligand(s), PD-1 can trigger apoptosis of antigen-specific (CD4+ or CD8+) T cells.

In addition to dual-tumor mice, single-tumor mice were examined as well. 3 groups of Immunocompetent male C57BL/6 mice were inoculated SC in the right flank with 5 x 10^5^ MC38 tumor cells. These single-tumor groups all received peritumoral biopolymer injections (100 μL/mouse). One hour later, group 1 (n=5) was IV administered saline control (QD x 5 days), and groups 2 and 3 (n=10) were IV administered SQP33 protodrug (28.6 mg/kg Dox Eq QD x 5 days; cumulative dose of 143 mg/kg Dox Eq). Group 3 also received TLR9a as an intratumorally after the last SQP33 dose at 25 μg per mouse. On Day 38, all remaining animals were treated with a second cycle of SQ3370 – 100-μL intratumoral SQL70 biopolymer injection followed by 5 daily doses (1 dose per day) of SQP33 protodrug – this time at 11.9 mg/kg/dose Dox Eq (59.3 mg/kg/cycle Dox Eq). Mice from group 3 also received a second dose of TLR9a at 25 ug per mouse. On Day 70, one mouse trending toward complete response (group 2), and 5 mice (group 3) were re-challenged with 5 x 10^5^ MC38 tumor cells inoculated SC at the left flank. A control group of 5 naïve mice were also inoculated with 5 x 10^5^ MC38 tumor cells on the same day.

### SQL70 Biopolymer Preparation

Synthesis of SQL70 biopolymer follows an optimized 3-step process that includes: (i) condensing the sodium salt of HA with a tetrazine derivative (3-methyl-1,2,4,5-tetrazine-6-phenylmethanamine HCl) in the presence of excess carbodiimide activating agent, (ii) purification using tangential flow filtration (TFF) to remove low molecular weight starting materials, reagents, and by-products, and (iii) sterilization by ultrafiltration. The derivatization reaction occurs in N-morpholinoethanesulfonic acid (MES) buffered aqueous solvent with activation of the HA carboxylic acid group by N-hydroxysulfosuccinimide and N,N-(dimethylaminopropyl)-N-ethyl carbodiimide HCl.

### SQP33 Protodrug Preparation

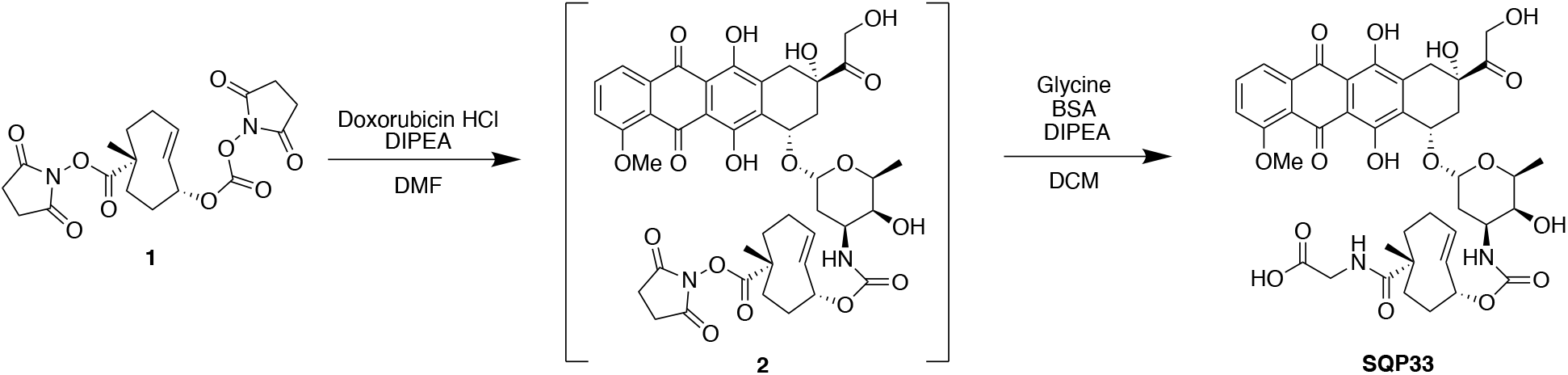

Compound **1** was synthesized using previously reported methods^5^. Compound **1** (15 g, 37 mmol) was suspended in 100 mL anhydrous DMF followed by the addition of doxorubicin hydrochloride (17.84 g, 30.76 mmol) and diisopropylethylamine (16 mL, 92 mmol). The suspension was stirred at rt for 3 h protected from light. Simultaneously, a suspension of glycine (23.1 g, 308 mmol), *N,O*-Bis(trimethylsilyl)acetamide (37.6 mL, 154 mmol), and diisopropylethylamine (107 mL, 615 mmol) in 440 mL anhydrous DCM was stirred at rt for 3 h. The DMF suspension was added to the DCM suspension in one portion and stirred at rt for 20 h. The reaction mixture was cooled to 0 °C, quenched with 1 L of ACN:water:formic acid (35:65:0.1), and brought to pH 3 with formic acid. DCM was removed under reduced pressure. Aliquots of 100 mL of the remaining solution were then purified by reverse phase prep-HPLC (step gradient ACN:water:formic acid 35:65:0.1 to 60:40:0.1, followed by 100% MeOH wash). Fractions were pooled and concentrated under reduced pressure to afford 25 g of orange solid. Aliquots of 0.8 g were dissolved in DCM:MeOH 94:6, and further purified by normal phase prep-HPLC (elution with DCM:MeOH 94:6). Fractions were pooled and concentrated under reduced pressure to afford SQP33 as an orange powder (15.46 g, 19.07 mmol, 62% yield). ESI+ [M+Na]: expected 833.27, observed 833.32. ESI-[M-H]: expected 809.28, observed 809.38.

### SQP33 Protodrug Solubility

Solid SQP33 protodrug was added to 1.0 mL 50 mM phosphate buffer at pH 7.0 until some solid remained undissolved and the saturation was reached. SQP33 protodrug dissolution significantly acidified the solution. The pH of the resulting suspension was then adjusted with small amount of concentrated sodium hydroxide solution (5 M) back to the target of 7.0. The two processes alternated until the vehicle was saturated (solid visible) at the target pH (± 0.5 pH unit). The suspension was then tumbled. The concentration of SQP33 protodrug was measured by HPLC of the supernatant at 24 h. The solubility of SQP33 protodrug was measured as 71.8 mg/mL. Under a similar experiment, the solubility of TCO-Dox (first generation) was <0.001 mg/mL.

### MTT assay

MC38 cells were seeded in 96-well microplates and exposed to SQP33 protodrug, Dox for 3 days using the The CellTiter 96^®^ AQueous One Solution Cell Proliferation Assay, a colorimetric method for determining the number of viable cells in proliferation or cytotoxicity assays. The assay reagent contains a tetrazolium compound [3-(4,5-dimethylthiazol-2-yl)-5-(3-carboxymethoxy-phenyl)-2-(4-sulfophenyl)-2H-tetrazolium, inner salt] and an electron coupling reagent (phenazine ethosulfate) (MTT assay). The quantity of formazan product as measured by absorbance at 490 nm is directly proportional to the number of live cells.

### Statistical Analysis

All the data was analyzed using GraphPad Prism 5. For our analysis, *p* < 0.05 was considered to be statistically significant. All results are expressed as means ± SEM or SD as indicated.

## Supporting Information

Figure S1 – MTT data from Max with MC38 cells.

Figure S2 – Body Weight data as a measure of toxicity.

Figure S3 – Rechallenge w/ or w/o TLR

**Supplemental Figure S1:**
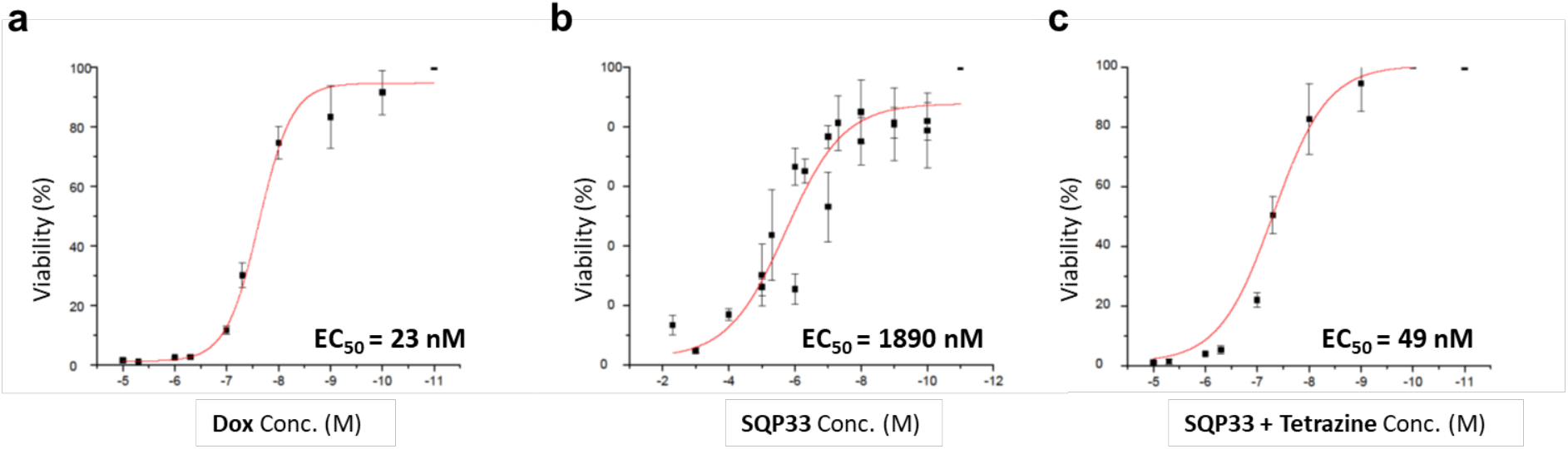
Cell Viability of MC38 Tumor Cell-line Was Assessed Using an MTT Assay. Cytotoxic effects of a) Dox, b) SQP33 protodrug, and c) SQP33 protodrug in the presence of tetrazine were compared were compared in an MC38 cell line after incubation for 3 days. Samples were run in triplicate. Percent viability is shown for increasing concentrations of each drug. EC50 is the concentration of a drug that gives half-maximal response. Without activation by tetrazine, the EC50 of SQP33 protodrug was 82-fold lower (b) than with tetrazine (c).

**Supplemental Figure S2:**
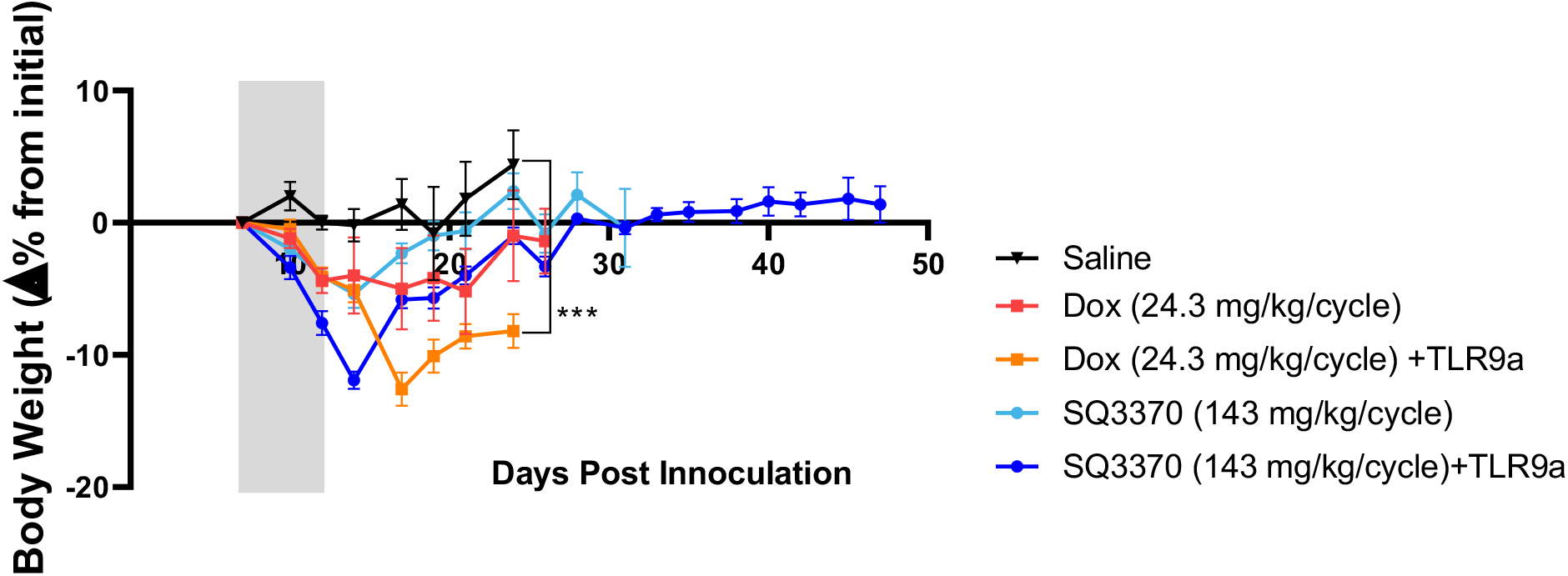
Changes in Body Weight in Tumor Bearing Mice after Treatment. Body weights as percent change from initial during the start of treatment shown for saline, Dox, Dox+TLR9a, SQ3370, and SQ3370+TLR9a treated groups, respectively. Gray bars represent the treatment duration. Data points without errors bars occurred when the standard error was smaller than the symbol used to represent the treatment condition. Curves stopped after 1 or more mice in that group died or were sacrificed when tumor volume reached 2000 mm3. Gray bars represent the SQ3370 treatment duration. Statistical significance in curves was determined by comparison to saline group using a one-way ANOVA with Tukey’s multiple comparisons for each day. *p<0.05; **p<0.01;***p<0.001.

**Supplemental Figure S3:**
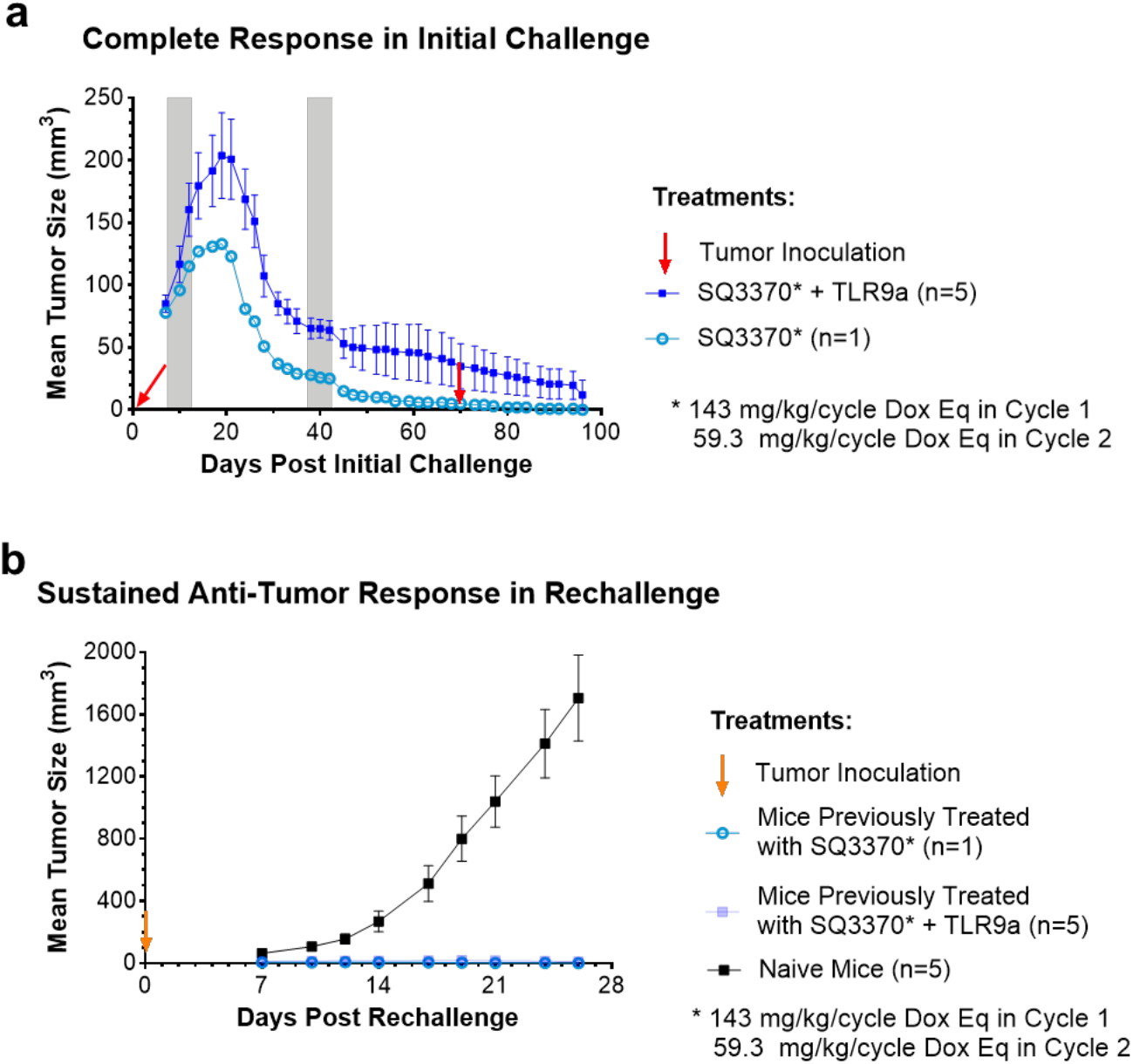
Sustained Immune Response After Tumor Re-challenge in SQ3370-Treated Mice. a) Immunocompetent C57BL/6 mice were inoculated with mouse MC38 tumor cells and treated with SQ3370 for 2 cycles at 143 mg/kg/cycle Dox Eq in Cycle 1 and 59.3 mg/kg/cycle Dox Eq in Cycle 2, with or without TLR9a in Cycle 2. Mice were re-challenged with MC38 tumors on Day 70 and each treatment group was compared with a control group of naïve mice also inoculated on the same day. b) Initial challenge and c) rechallenge tumor growth curves are shown for each treatment group. Gray bars represent the duration of treatment in each cycle. Tumor growth curves show mean ± SEM.

